# Topographic gradients define the projection patterns of the claustrum core and shell in mice

**DOI:** 10.1101/2020.09.11.293381

**Authors:** Brian A. Marriott, Alison D. Do, Ryan Zahacy, Jesse Jackson

**Author notes:** **CORRESPONDENCE**. 7-22 Medical Sciences Building, University of Alberta. Edmonton, Alberta, Canada, T6G 2H7.

## Abstract

The claustrum is densely connected to the cortex and participates in brain functions such as attention and sleep. Although some studies have reported the widely divergent organization of claustrum projections, others describe parallel claustrocortical connections to different cortical regions. Therefore, the details underlying how claustrum neurons broadcast information to cortical networks remain incompletely understood. Using multicolor retrograde tracing we determined the density, topography, and co-projection pattern of fourteen claustrocortical pathways, in mice. We spatially registered these pathways to a common coordinate space and found that the claustrocortical system is topographically organized as a series of overlapping spatial modules, continuously distributed across the dorsoventral claustrum axis. The claustrum core projects predominantly to frontal-midline cortical regions, whereas the dorsal and ventral shell project to the cortical motor system and temporal lobe, respectively. Anatomically connected cortical regions receive common input from a subset of claustrum neurons shared by neighboring modules, whereas spatially separated regions of cortex are innervated by different claustrum modules. Therefore, each output module exhibits a unique position within the claustrum and overlaps substantially with other modules projecting to functionally related cortical regions. Claustrum inhibitory cells containing parvalbumin, somatostatin, and neuropeptide Y also show unique topographical distributions, suggesting different output modules are controlled by distinct inhibitory circuit motifs. The topographic organization of excitatory and inhibitory cell types may enable parallel claustrum outputs to independently coordinate distinct cortical networks.

## INTRODUCTION

The claustrum communicates most prominently with the cortex, and has one of the densest connectivity profiles per unit volume in the forebrain (Atlan et al., 2017; Edelstein & Denaro, 2004; Milardi et al., 2015; Torgerson et al., 2015; Q. Wang et al., 2017; Zingg et al., 2014). Excitatory outputs from the claustrum activate cortical inhibitory interneurons leading to feedforward inhibition of cortical activity (Atlan et al., 2018; Cortimiglia et al., 1991; Jackson et al., 2018; Narikiyo et al., 2020). Recent evidence has shown that the claustrum participates in a diverse array of functions including sleep, attention, and memory (Atlan et al., 2018; Goll et al., 2015; Liu et al., 2019; Narikiyo et al., 2020; Norimoto et al., 2020; Renouard et al., 2015; White et al., 2020). Considering the diversity of proposed claustrum functions and the widespread connectivity with the cortex, we sought to determine the organization of claustrocortical projections.

Previous anatomical studies show that a subset of claustrum neurons send axon collaterals throughout the entire cortical axis (Atlan et al., 2018; Q. Wang et al., 2017; Y. Wang et al., 2019; Zingg et al., 2014, 2018), and activation of the claustrum can evoke changes in cortical activity across widely distributed regions of cortical space (Narikiyo et al., 2020). These data have inspired the proposal that the claustrum could serve to broadly coordinate disparate regions of the cortex. However, other retrograde tracing data suggest claustrum connections to the cortex are compartmentalized – linking separate populations of claustrum neurons with different cortical regions (Chia et al., 2020; Gattass et al., 2014; Macchi et al., 1983; Minciacchi et al., 1985; Sadowski et al., 1997; Smith et al., 2019; Watson et al., 2017; White et al., 2017). This latter hypothesis has received strong support, as all species studied to date show separate sets of claustrum neurons projecting to distinct areas of sensory cortex (LeVay & Sherk, 1981; Olson & Graybiel, 1980; Remedios et al., 2010; Smith et al., 2012). In many cases, these different output streams are topographically organized (Gattass et al., 2014; Li et al., 1986; Minciacchi et al., 1985; Pearson et al., 1982; Reser et al., 2017; Watson et al., 2017; Witter et al., 1988), whereas in others (particularly in rodents), a lack of topography is noted (Chia et al., 2020; White et al., 2017; Zingg et al., 2018). Discrepancies between studies likely arise from different claustrum projections being measured, and species-specific differences in claustrocortical organization (Binks et al., 2019; Orman et al., 2017; Pham et al., 2019; Smith et al., 2019). As mice provide a powerful model system for studying cell types, circuits, and behavior, a comprehensive understanding of the mouse claustrocortical system is required.

Here, we systematically measured the density, spatial organization, and collateralization of claustrocortical projections to different cortical regions. In doing so, we found a continuum of overlapping claustrocortical modules organized primarily along the dorsoventral axis. This topographical organization ensures that spatially distant and weakly connected cortical regions receive inputs from independent claustrum populations, while neighboring and connected cortical regions receive common claustrum inputs. Coupled with this output topography, we found interneurons containing somatostatin and neuropeptide – Y were spatially organized and exhibit a particularly dense labelling in the claustrum relative to surrounding cortical regions. Knowledge of these anatomical motifs will guide future experiments aimed at determining if distinct claustrum populations have unique roles in cognition.

## MATERIALS AND METHODS

All procedures were performed according to the Canadian Council on Animal Care Guidelines and were approved by the University of Alberta Animal Care and Use Committee (AUP2711). Male and female C57BL/6 mice, between 60-180 days old, were used for all experiments. Mice were group housed in a temperature-controlled environment on a reverse 12-hour light-dark cycle. NPY-hrGFP mice (van den Pol et al., 2009) were obtained from Jackson labs (RRID: IMSR_JAX:008069).

### Tracer injection

Mice were administered carprofen via ad-libitum water 24 hours prior to surgery, and for 72 hours after surgery to achieve a dose of 5mg/kg. For surgery, mice were initially anesthetized using 4% isoflurane and maintained at 1.0-2.5%. Mice were secured in a stereotaxic frame, with body temperature maintained through an electric heating pad set at 37°C. Local anesthetic (bupivacaine) was applied locally under the scalp, and an incision along midline was made to access bregma and all injection sites. The skin was moved back from the intended injection sites using sterile swabs and kept moist during surgery with sterile 0.9% saline. The skull was leveled between bregma and lambda. Craniotomies were marked and manually drilled using a 400μm dental drill bit according to stereotaxic coordinates (**Table 1**), and the dorsoventral measurements made from brain surface. The left hemisphere was used for all injections unless otherwise stated. Pulled pipettes (10-20μm in diameter) were back filled with mineral oil and loaded with tracers. All injections were made using pressure injection. The glass pipette was lowered into the injection site at 1mm per minute, and 150-200nL of each tracer was injected at 50-100nL/minute. The pipette was allowed to rest for 10 minutes after injecting before removal. Fast blue (Bentivoglio et al., 1980; Kuypers et al., 1980) (Polysciences, Pennsylvania) was prepared by dissolving 1mg of powder in 30μL 1X phosphate buffered saline (PBS) and 1.5μL of dimethyl sulfoxide. The solution was warmed and agitated to fully dissolve and was stored at 4°C in 3μL aliquots. Cholera Toxin subunit-B (Luppi et al., 1995) with a Alexa Fluor-647 (AF-647) conjugate (ThermoFisher, catalog number C34778) was prepared by dissolving 100μg in 20μL PBS, agitated to dissolve, stored at 4°C, and gently vortexed before injection. Retrograde adeno associated viruses encoding green fluorescent protein (Addgene, Catalog number 50465-AAVrg) or tdtomato (Addgene, Catalog number 59462-AAVrg) were obtained from Addgene, and aliquoted (3μL) and stored at −80°C. Prior to surgery an aliquot was thawed on ice. The skin was sutured after completing all injections and sealed with vetbond (3M). Mice were returned to fresh cages upon regaining consciousness.

**Table 1:** The cell counts and the percentage of neurons in the core and shell across the rostrocaudal claustrum axis.

| Region | ALM | MOs | PL | MOp | ACA | aRSP | SSbfd | iRSP | AUDp | VISp | pRSP | ENTl | pSUB | ENTm |
| --- | --- | --- | --- | --- | --- | --- | --- | --- | --- | --- | --- | --- | --- | --- |
| **Coordinates**<br>A-P<br>M-L<br>D-V (mm from<br>brain surface) | 2.5<br>1.5<br>-0.8 | 1.8<br>0.7<br>-0.5 | 1.7<br>0.5<br>-1.0 | 1.0<br>1.7<br>-0.8 | 0.5<br>0.3<br>-0.8 | -1.0<br>0.5<br>-0.5 | -1.0<br>2.8<br>-0.8 | -1.5<br>0.3<br>-0.5 | -2.5<br>3.8<br>-1.0 | -2.9<br>2.5<br>-0.5 | -3.0<br>0.5<br>-3.0 | -3.0<br>3.8<br>-3.0 | -4.0<br>2.0<br>-1.5 | -4.6<br>3.0<br>-2.7 |
| **Tracer** | rGFP | FB | FB,<br>rGFP,<br>CTB | rTD | rGFP | CTB,<br>rTD | FB | All | rGFP | FB,<br>rTD | rGFP,<br>FB | rGFP | FB | rTD |
| **Rostral cell**<br>**counts** | 116±24 | 125±11 | 171±74 | 90±14 | 74±3 | 117±47 | 23±4 | 73±23 | 36±17 | 44±17 | 32±8 | 91±45 | 51±13 | 110±16 |
| **Rostral.**<br>**Dorsal shell %** | 70±5 | 24±4 | 25±8 | 62±6 | 10±4 | 17±10 | 43±5 | 5±2 | 34±10 | 23±11 | 6±6 | 11±5 | 9±4 | 16±5 |
| **Rostral**<br>**Core %** | 23±4 | 57±3 | 51±5 | 31±3 | 47±9 | 76±14 | 50±8 | 90±0 | 43±12 | 51±16 | 80±3 | 31±9 | 65±15 | 40±12 |
| **Rostral**<br>**Ventral shell %** | 7±1 | 19±7 | 24±8 | 7±3 | 43±5 | 7±5 | 7±3 | 5±2 | 23±7 | 26±10 | 14±3 | 58±7 | 26±12 | 44±12 |
| **Intermediate**<br>**Neurons counts** | 62±12 | 80±22 | 91±36 | 60±15 | 45±6 | 88±39 | 16±5 | 46±16 | 26±3 | 31±5 | 36±10 | 78±42 | 52±15 | 59±20 |
| **Intermediate**<br>**dorsal shell %** | 87±4 | 40±6 | 20±4 | 81±8 | 14±2 | 17±11 | 64±7 | 5±1 | 40±6 | 16±6 | 4±2 | 19±8 | 10±3 | 13±5 |
| **Intermediate**<br>**Core %** | 8±2 | 42±2 | 55±4 | 13±4 | 45±13 | 73±17 | 30±7 | 90±0 | 37±7 | 57±13 | 82±6 | 19±9 | 68±12 | 44±11 |
| **Intermediate**<br>**Ventral shell %** | 5±2 | 18±6 | 25±5 | 6±4 | 41±10 | 10±6 | 6±2 | 5±1 | 23±10 | 27±9 | 14±5 | 62±12 | 22±8 | 43±10 |
| **Caudal neuron**<br>**counts** | 30±7 | 41±10 | 44±26 | 27±5 | 25±2 | 38±5 | 9±2 | 29±10 | 11±5 | 20±3 | 26±3 | 67±36 | 30±5 | 40±11 |
| **Caudal**<br>**Dorsal shell %** | 76±8 | 39±6 | 30±10 | 79±5 | 21±10 | 24±17 | 65±14 | 5±1 | 39±19 | 20±7 | 7±4 | 15±7 | 13±6 | 17±4 |
| **Caudal**<br>**Core %** | 11±5 | 36±5 | 41±12 | 13±5 | 30±11 | 61±29 | 25±9 | 90±0 | 34±19 | 55±12 | 71±18 | 7±3 | 55±18 | 33±9 |
| **Caudal**<br>**Ventral shell %** | 13±4 | 25±9 | 29±2 | 8±2 | 49±1 | 15±12 | 10±6 | 5±1 | 27±3 | 25±13 | 21±20 | 78±9 | 32±14 | 50±7 |

### Perfusion and tissue sectioning

Mice were deeply anesthetized and transcardially perfused 2-3 weeks after injections with ice cold PBS, followed by 4% paraformaldehyde (PFA) in PBS. Brains were extracted and post-fixed in 4% PFA for 24-48 hours and stored in PBS at 4°C until sectioning. Brains were mounted in 2% agarose and sectioned at 50μm using a vibratome (Leica VT1000s, Germany). Coronal sections were used for all brains. The entire brain was sectioned, and every second slice mounted on glass slides and sealed with coverslips using Prolong Gold (ThermoFisher). Slides were kept at 4°C until imaging.

### Immunohistochemistry

Mice were perfused and coronal sections obtained as above. Slices were first washed with 1X PBS (3 x 10 min) and then blocked using 2% bovine serum albumin in PBST (0.4% Triton X-100 in 1X PBS) for 2 hours at room temperature (RT). Sections were incubated with primary antibody rat anti-somatostatin (1:250, Millipore cat. No. MAB354, RRID:AB_2255365) at RT for 24 hours and then 4°C for 42 hours. For parvalbumin (PV) immunohistochemistry, slices were incubated in goat anti-Paravalbumin (1:2000, Swant, RRID:AB_10000345) for 18 h at 4°C. The slices were then washed with 0.1% PBST (3 x 10 min) followed by incubation with fluorophore-conjugated secondary antibodies: donkey anti-Rat Dylight 488 or 647 (1:500, Invitrogen) and donkey anti-Goat Alexa 647 (1:500, Invitrogen) at RT for 4 hours. After washing with 0.1% PBST (3 x 10 min) and then 1X PBS (3 x 10 min), slices were mounted onto slides and cover slipped. Confocal images were obtained on a Leica SP5 or SP8 using a 10x, 20x, or 25x objectives as described below.

### Imaging

Injection site images were taken on a widefield Zeiss AxioObserver.Z1 (Zeiss, Germany) with DAPI, EGFP, CY3, and CY5 filter cubes excited by 350nm, 488nm, 543nm, and 633nm LEDs respectively. Images used for neuron counts and co-localization analysis were taken on a Leica DMI6000B SP8 (Leica, Germany) confocal microscope with a 10x 0.4NA or 25x 1.0NA objective, using a 405nm laser, and a white light laser set at 488nm, 543nm, and 633nm with the acousto-optical emission filtering set using Leica defaults for DAPI, EGFP, tdTomato, and AF-647. Fast blue was detected with a conventional PMT, and EGFP, tdTomato, and AF-647 were detected using Hybrid Detectors. Laser intensity and detector settings were adjusted for each brain to optimize brightness and contrast for each channel. Six slices from each brain were imaged for analysis. This included two rostral claustrum slices separated by 200μm, two intermediate claustrum slices separated by 200μm, and two caudal claustrum slices separated by 200μm. Both the ipsilateral and contralateral claustrum (relative to injections) were imaged in each brain. Images were taken at 2048×2048 pixels, accumulation = 2x, bidirectional x, pinhole set to 1 airy unit, and a z-stack of 4 images over a 12μm volume were taken. Each scan was set to image fast blue and AF-647 simultaneously, with EGFP and tdTomato imaged sequentially. Images were loaded into FIJI and converted to maximum intensity z-projection for analysis. We found that retrograde labelling of neurons in the claustrum was spatially sparse enough in the z-imaging plane to enable analysis using the maximum intensity projection (over this small volume), as the manual assessment of co-localization using multiple z-axis imaging planes or the maximum intensity projection yielded the same rate of co-projections on a subset of images analyzed.

### Analysis

Only pathways where the injection site was confirmed to reside in the target region were used for analysis. Before quantification, images were rotated (if necessary) such that the dorsoventral axis was vertical and parallel to the y-axis of each image. Images were quantified in Matlab by manually counting and recording the location of neurons in each imaging channel for each image using cursor clicks that stored the x-y coordinate for each neuron within the image. After identifying all neurons in each channel, the determination of co-projections was performed by finding pairs of neurons (across channels) that were within 50μm of each other, and these neurons were replotted for manual inspection of co-labelling at high magnification. This was repeated for all six pairwise comparisons for four channel images. The x-y coordinates of all neurons in each channel and image were then used for a second round of manual co-labelling measurements for triple and quadruple label expressing neurons. All neurons x-y coordinates were registered to the CLA_RSP_ pathway as all brains had retrograde tracers in the same RSP coordinate. For registration, the centroid of the CLA_RSP_ neurons was calculated and used for centering all other neurons in the x-y direction. Therefore, each neuron was assigned a new, normalized, x-y coordinate representing the distance from this CLA_RSP_ centroid. The perimeter of CLA_RSP_ neurons was determined by using the perimeter function in Matlab, using the closest 90% of the neurons to the CLA_RSP_ centroid. This polygon defined the claustrum core. Neurons located outside and dorsal to the CLA_RSP_ core were defined as being in the dorsal shell, whereas neurons located outside and ventral were defined as ventral shell. We did not include a dorsal or ventral limit on the extent of the dorsal or ventral shell. Instead we used the histograms and density plots to display where claustrocortical cells were located. The spatial density of all claustrocortical projections was generated using 30μm x 30μm bins, and all cells in the imaging field of view were included in the analysis. The spatial density of claustrocortical projections was then measured in both the mediolateral and dorsoventral axes and compared with the CLA_RSP_ reference population. From the spatial density maps, the outline of each claustrocortical projection was made using Otsu’s method (Matlab), whereby a spatial threshold is determined that minimizes the intra-class (within boundary and outside boundary) variance of the values in each bin. Co-labelling between two pathways was calculated by dividing the number of double labelled neurons by the sum of the all labelled neurons across the two regions minus the double labelled neurons. For example, the proportion of neurons projecting to both regions A and B = AB / (A + B – AB). Four color tracing yielded fifteen different types of labelling patterns that comprised single, double, triple, and quadruple labelling. The number of single, double, triple, and quadruple labelled neurons were summed and represented in histogram form and pie charts.

### Cortical connectivity estimation

Data were obtained from Supplemental Table 3 from Oh et al., 2014. Data in the matrix table reflect the projection strength values extracted from anterograde fluorescence tracing between cortical regions. For each pair of cortical regions, the table described a source (injection site), target (post synaptic region), and the adjusted intensity of axon labelling (see Oh et al., for details). We averaged the connectivity estimate across both directions of each pair of cortical regions to obtain a single value reflecting the relative connectivity strength between regions. The table contains source – target connectivity density information for most cortical regions. However, data for our ALM coordinate and different rostrocaudal levels of the RSP were not differentiated in this data. Therefore, for our cortical connectivity analysis, the RSP was considered a single structure, and ALM was not included. Consequently, nineteen pairs of cortical regions were compared, rather than the original twenty-seven.

### Statistics

The mean and standard deviation (across mice or slices) are shown in all Figures, unless otherwise stated. Pairwise t-tests or Wilcoxon rank sum tests were used and corrected for multiple comparisons with the Bonferroni correction. P-values of < 0.05 were deemed statistically significant.

## RESULTS

The claustrum was studied using a series of coronal brain sections from across the rostrocaudal axis (**Figure 1a**), giving high spatial resolution in the dorsoventral and mediolateral axes (see Materials and Methods). First, we required a consistent anatomical landmark to spatially register claustrum neurons across experiments. Parvalbumin (PV) neuropil labelling, and retrograde tracing from the retrosplenial cortex (RSP) have both been used to topographically locate the claustrum (Dillingham et al., 2019; Druga et al., 1993; Mathur et al., 2009; Q. Wang et al., 2017; White et al., 2017; Zingg et al., 2018). Comparing these two markers in dorsoventral, mediolateral, and rostrocaudal axes, showed a highly correlated spatial overlap, indicating that both methods identify a common region of the claustrum (**Figure 1a-c**). Therefore, we chose to use the claustrum -> RSP (CLA_RSP_) pathway to align the retrograde labelling from other cortical regions. The center of mass of CLA_RSP_ neuron labelling was defined as the center of the claustrum, and all retrograde labelled neurons in each coronal brain section were spatially re-aligned to this common coordinate space as shown in **Figure 1d-e** (see Methods and Materials). A polygon defined by the perimeter of CLA_RSP_ neurons was used to demarcate the claustrum core. This approach ensured that the boundaries of the claustrum and the location of each neuron was determined objectively and without bias. Retrograde tracers were deposited in three to four cortical regions within each brain, for a total of fourteen regions injected across all experiments. For each brain, one tracer was injected into the RSP at an intermediate location along the rostrocaudal axis (−1.5mm from bregma), and all other tracers deposited into anatomically distinct areas of the cortex (**Table 1**). The full range of cortical injection sites included the anterior lateral motor cortex (ALM), primary motor cortex (MOp), secondary motor cortex (MOs), prelimbic cortex (PL), rostral retrosplenial cortex (rRSP), intermediate retrosplenial cortex (iRSP), caudal retrosplenial cortex (cRSP), somatosensory barrel cortex (SSbfd), primary auditory cortex (AUDp), primary visual cortex (VISp), anterior cingulate (ACA), post-subiculum (pSUB), medial entorhinal cortex (ENTm), and lateral entorhinal cortex (ENTl) (**Table 1**). Averaging across all brains we found retrograde labelling from these cortical injection sites showed a considerable spatial spread, beyond the border defined by CLA_RSP_ and PV labelling (**Figure 1f**, middle). Thus, we adopted the term ‘core’ and ‘shell’ to provide coarse-grained classification of the spatial location of retrogradely labelled neurons (**Figure 1f,** right) in accordance with the core-shell nomenclature used previously (Atlan et al., 2017; Real et al., 2006).

**Figure 1.**
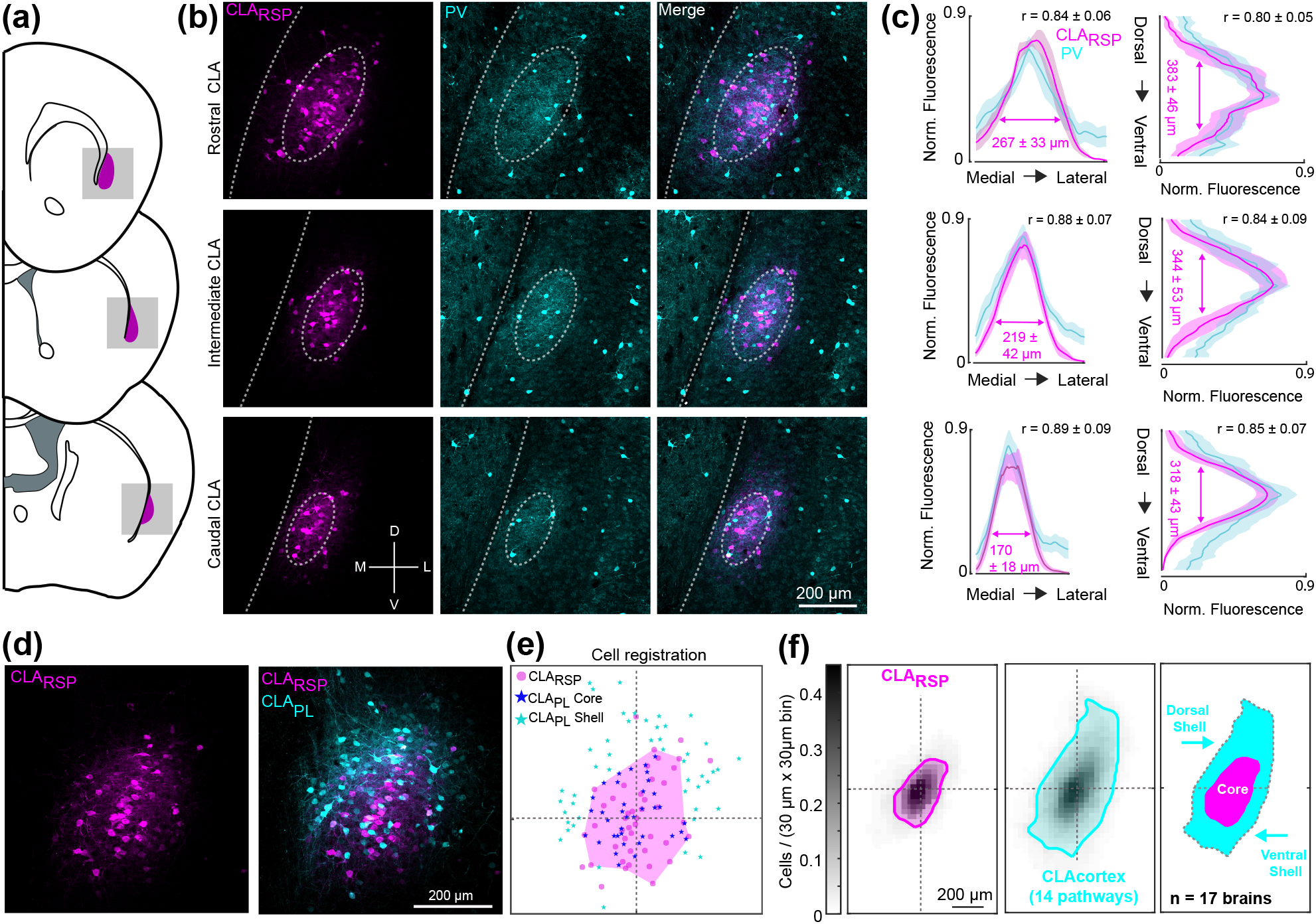
Spatial registration of claustrocortical neurons across brains. **(a**) Schematic coronal sections showing the rostral, intermediate, and caudal claustrum. (**b**) Examples of retrogradely labelled CLA_RSP_ neurons (magenta) and parvalbumin (PV, cyan) immunohistochemistry across the rostrocaudal axis. (**c**) The quantification of PV and CLA_RSP_ labelling in the rostral (top), intermediate (middle), and caudal (bottom) claustrum, in both mediolateral (left) and dorsoventral (right) axes. The correlation coefficient between PV and CLA_RSP_ is shown in the top right of each plot (n = 3 mice, 6 slices at each spatial location). (**d**) Two color retrograde tracing from the RSP and prelimbic cortex (PL). (**e**) The spatial registration of the CLA_PL_ pathway in this example image, using the CLA_RSP_ pathway as a reference. The magenta polygon outlines the spatial extent of CLA_RSP_ labelling (See Methods and Materials). Neurons inside/outside of the CLA_RSP_ region are classified as core/shell respectively. For each image, 10% of CLA_RSP_ neurons most distant from the CLA_RSP_ centroid were removed before calculating the claustrum core polygon, in order to reduce the effect of spatial outliers. (**f**) The average spatial density of CLA_RSP_ neurons (magenta, left), and the density of all fourteen claustrocortical pathways studied (cyan, middle). The overlay of the two plots shows regions classified as the dorsal shell, core, and ventral shell (far right). Otsu’s method (see Methods) was used to calculate the boundaries of the core and shell for these density plots.

The three tracer types included fast blue (FB), fluorescently tagged cholera toxin subunit-B (CTB-647), and two variants of AAV2-retro (AAV2-retro-tdtomato, and AAV2-retro-GFP) (Tervo et al., 2016). As our goal was to compare the number of claustrocortical neurons projecting to several cortical areas using different tracers, we first determined if each retrograde tracer showed comparable tracing efficacies. We found that a similar number of claustrum neurons were detected when injected into the RSP (Fast blue: 44±13; CTB: 46.7±14.7; AAVretro: 41.5±7.5 neurons/slice, **Figure 2a-b**), V1 (fast blue: 33.7±14.4; AAVretro: 33.6±2.4, **Figure 2c-e**) and M2 (Fast blue: 81.7±14.7; AAVretro: 67.9±6.5 neurons/slice) (**Figure 2d-f**). Likewise, different tracer combinations led to similar rates of co-projecting neurons detected in the claustrum. Therefore, these tracers have a similar efficacy, do not compete, and can be used for multicolor claustrocortical mapping in the same brain. Example injection site locations are shown in **Figure 3.**

**Figure 2:**
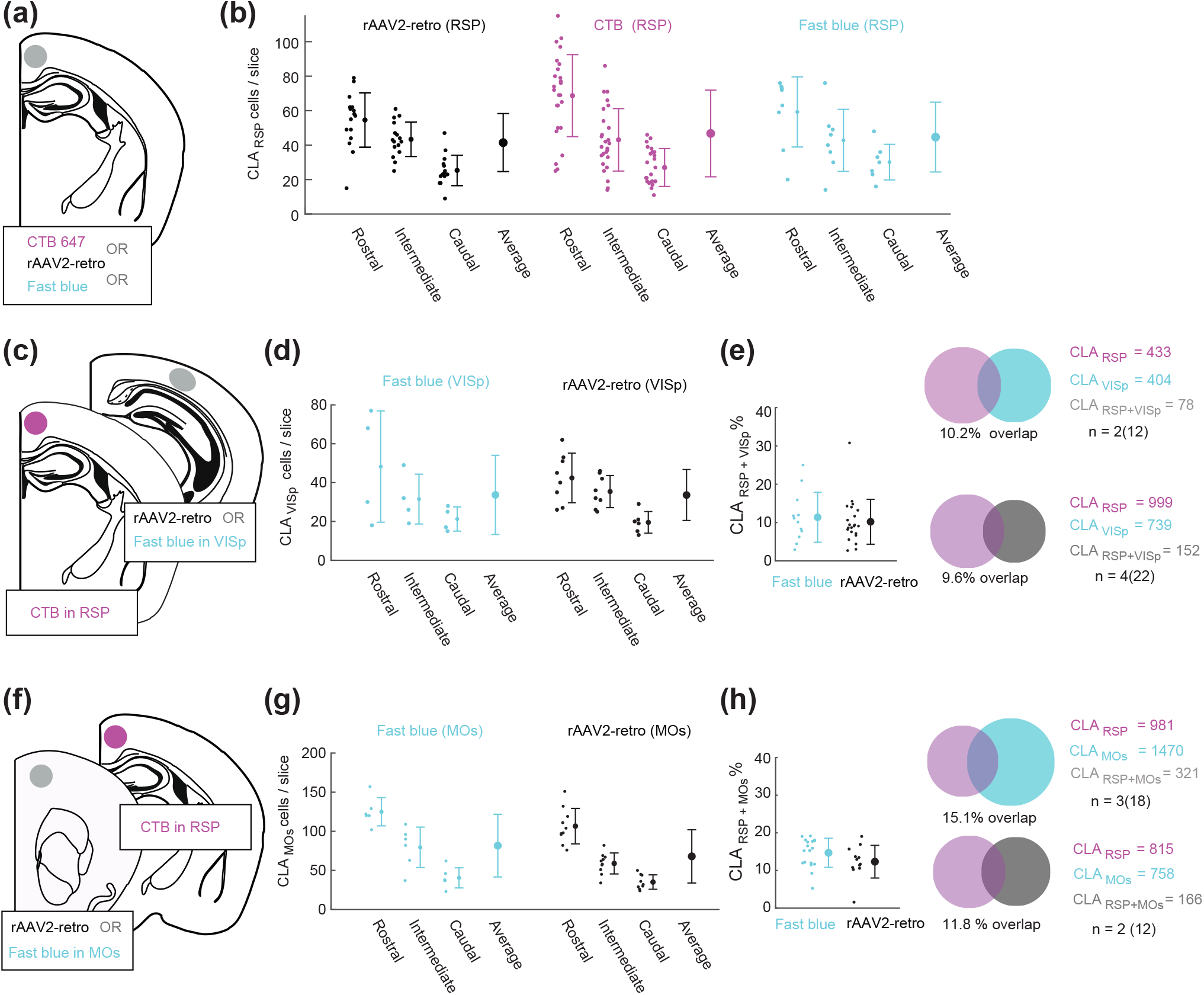
Comparing the efficacy of different retrograde tracers. (**a**) A schematic showing the injection of CTB647, rAAV2-retro-GFP (or rAAV2-retro-tdTomato), or fast blue into the intermediate retrosplenial cortex (iRSP). (**b**) The number of CLA_RSP_ neurons detected using each tracer type. The neuron counts were performed in the ipsilateral claustrum at rostral, intermediate, and caudal levels of the claustrum. Each point is from one slice and the mean and standard deviation are shown across all sections. No difference in the average number of neurons counted for each tracer were detected, and all tracers showed similar profiles of decreased claustrum labelling along the rostrocaudal axis. (**c**) A schematic depicting the injection of rAAV2 or fast blue into primary visual cortex (VISp) together with CTB-647 into the RSP. (**d**) The number of CLA_VISp_ neurons labelled with rAAV2 or fast blue was not significantly different. (**e**) The number of co-labelled claustrum neurons projecting to RSP and VISp was not significantly different between the experiments where fast blue was used (n = 2 mice, 12 slices), or rAAV2 was used (n = 4 mice, 22 slices). (**f-h)** The same as (**c-e**) except for the CLA_MOs_ pathway. There was no difference between the number of neurons labelled by fast blue or rAAV2 in the CLA_MOs_ pathway, and the percentage of co-labelled neurons was not significantly different between experiments with CLA_RSP_(CTB) + MOs (fast blue) or CLA_RSP_ (CTB) + MOs (rAAV2).

**Figure 3.**
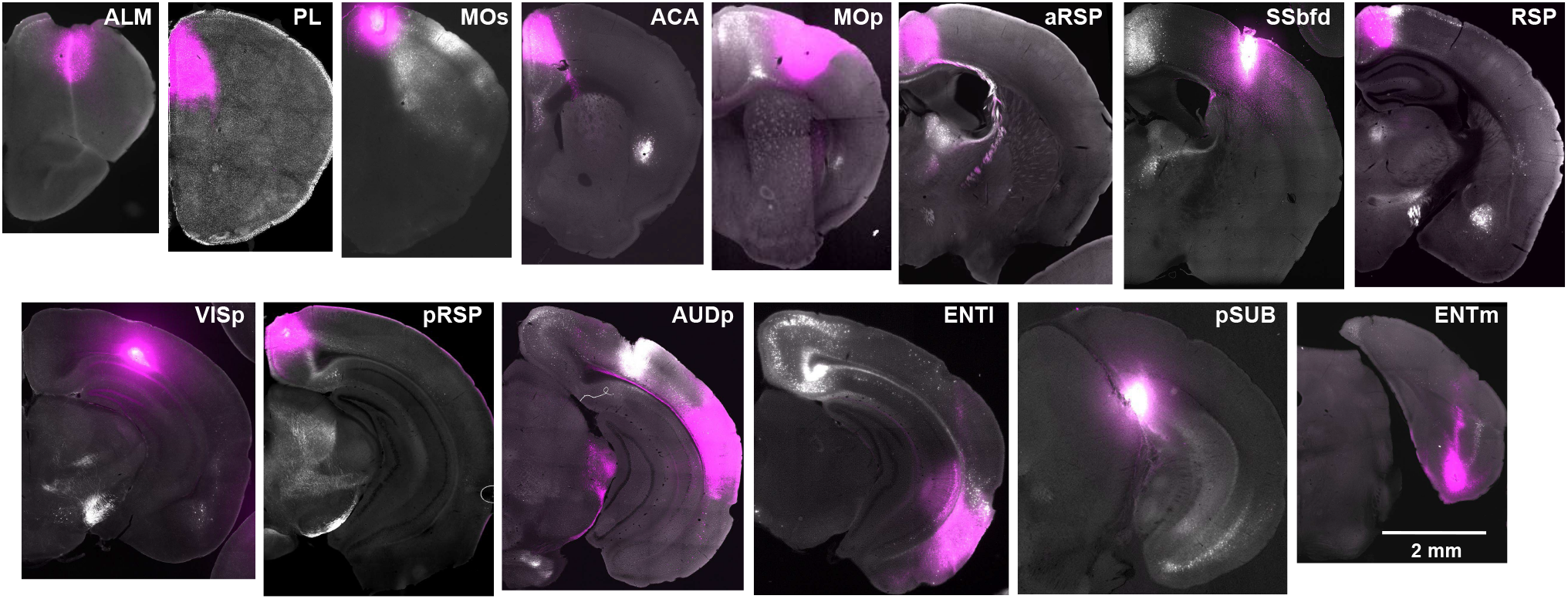
Example injection sites for retrograde tracing of claustrocortical projections. Coronal brain sections show the location of the retrograde tracer injections sites (magenta).

The topography of individual projections was assessed in the dorsoventral, mediolateral and rostrocaudal axis. Visualization of up to four different claustrocortical pathways in the same brain revealed that different pathways were differentially distributed across the dorsoventral claustrum axis (**Figure 4, Figure 5a-b**), and shifts in the mediolateral axis were attributed to the slight claustrum curvature dorsoventrally (**Figure 5c**). The projections to ALM and PL showed proportionally more labelling in the rostral claustrum, whereas projections to the pSUB, ENTm, and ENTl had more neurons in the caudal claustrum, relative to the reference CLA_RSP_ pathway (**Figure 5d-e, Table 1**). To quantify the dorsoventral topography more simply, we compared the proportion of claustrocortical neurons in the dorsal shell, core, and ventral shell for each cortical injection region. Projections to ALM, MOp, and SSbfd were significantly increased in the dorsal shell, (**Figure 5a-f**), whereas projections to the ACA, ENTm, and ENTl were biased toward the ventral shell. Claustrum projections to AUDp and PL were spread more equally between core and shell, and the projections to VISp and pSUB had a topography that most closely mirrored the CLA_RSP_ neurons, but with a shift toward the ventral shell (**Figure 4b and d, and Figure 5**). The distribution of core/shell neurons within each pathway was largely conserved across the rostrocaudal axis (**Table 1**). Retrograde labelling in the contralateral claustrum occurred at rate of 0-16% of that detected in the ipsilateral claustrum (**Figure 6a-c**), in accordance with the tracing data described previously (Q. Wang et al., 2017). These neurons were mainly found in the rostral pole of the claustrum and sent inputs to the contralateral ALM, MOp, PL, and MOs, whereas contralateral projecting cells were nearly absent in the case of injections into VISp, AUDp, or areas of the temporal lobe (**Figure 6c**). Therefore, we propose that each claustrocortical pathway comprises a unique topographical position within the claustrum, yet its boundaries overlap considerably with several other pathways.

**Figure 4.**
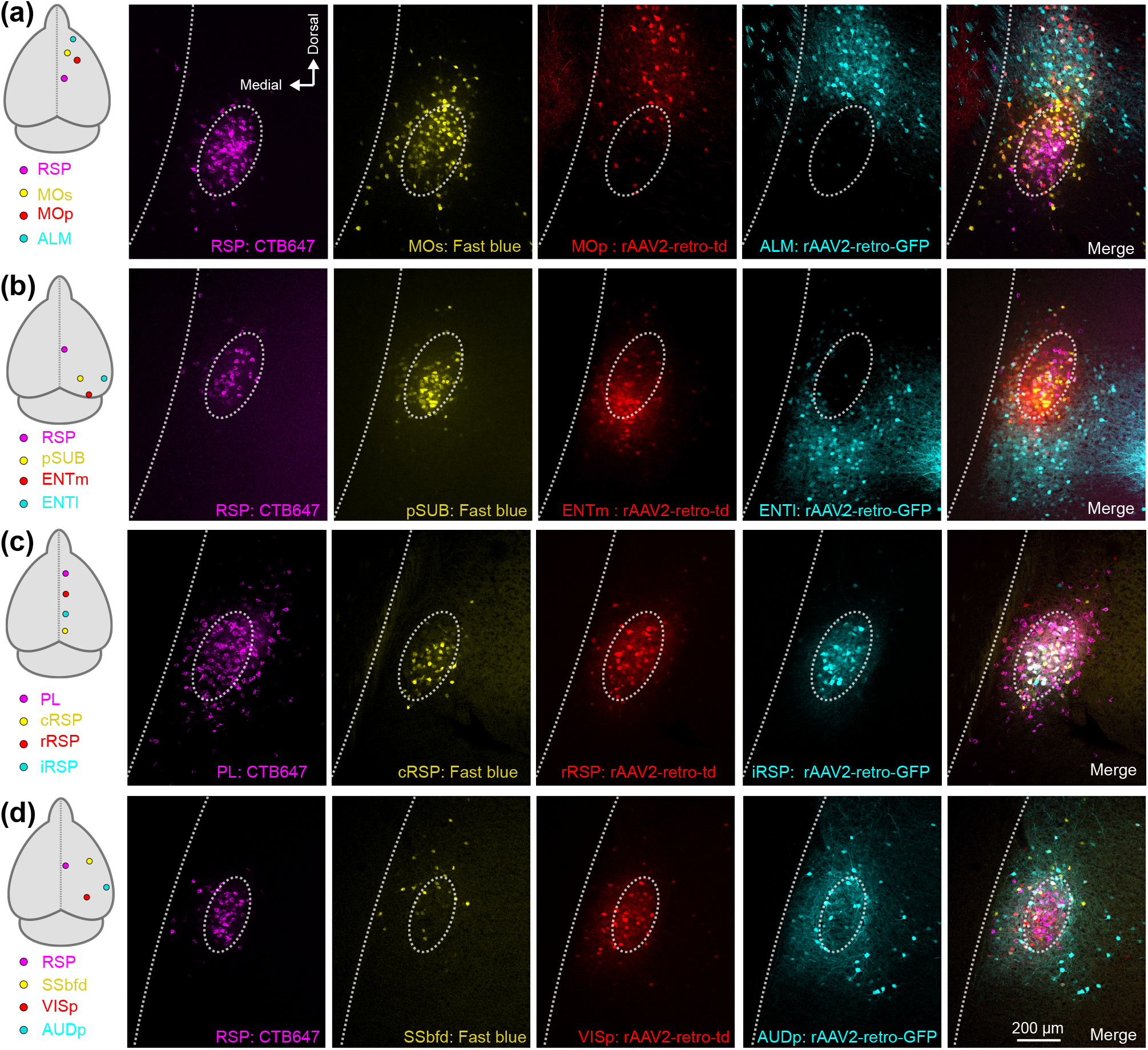
Diverse spatial domains of claustrocortical pathways. Four color tracing was performed in several sets of cortical injection configurations, four of which are shown (**a-d**). (**a**) Retrograde labelling in the claustrum following injections into the cortical regions indicated on the left (RSP, MOs, MOp, and ALM). The dashed oval is provided for visual alignment to the CLA_RSP_ pathway across all single channels and the merged image (far right). (**b-d)** The same as (**a**), for experiments with retrograde tracers targeting different areas of the temporal lobe (**b**), frontal-midline cortex (**c**), and sensory cortex (**d**). Data from other tracer combinations can be found in **Tables 1 and 2**.

**Figure 5.**
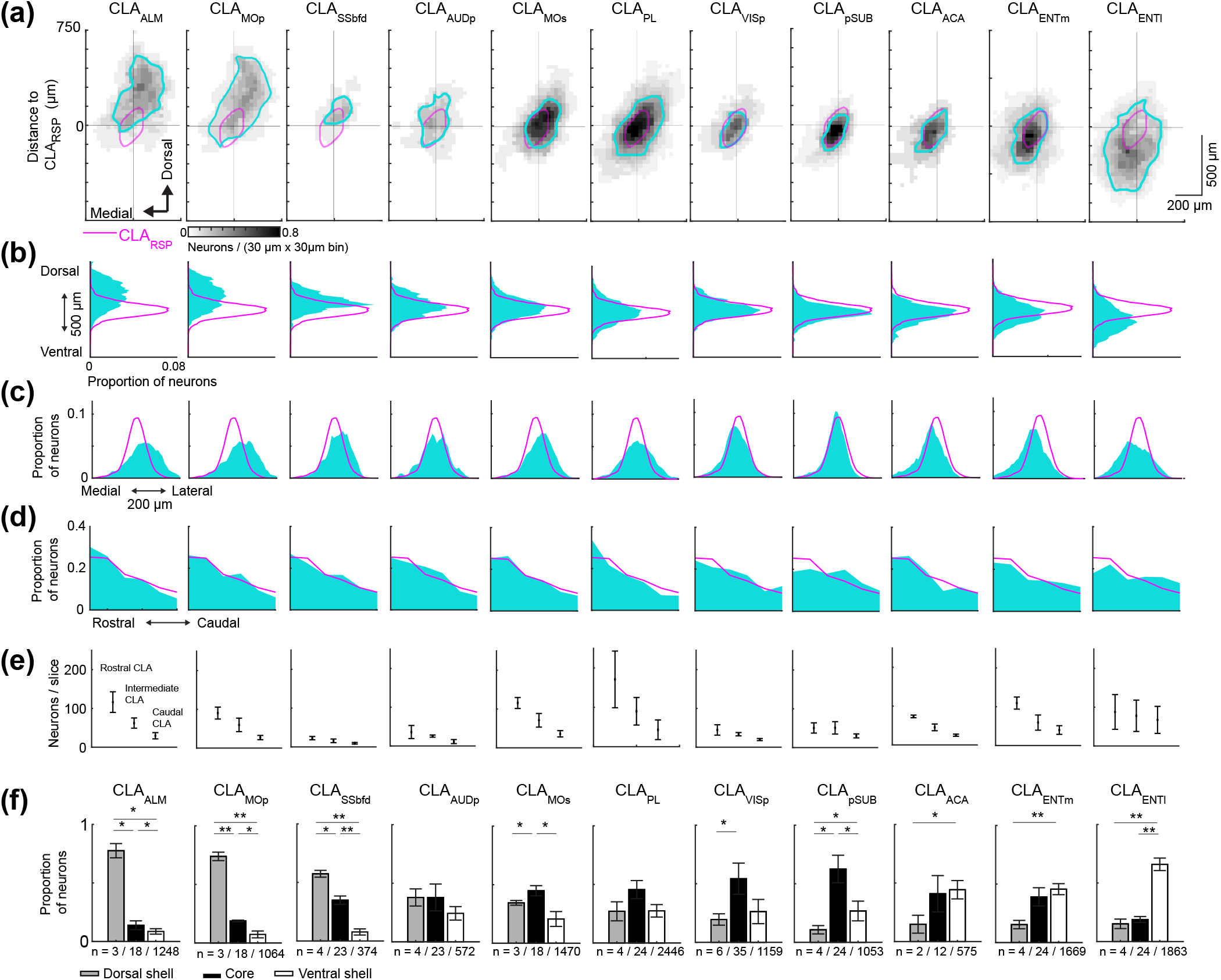
The topography of claustrocortical projections. (**a**) Spatial density maps of each claustrocortical pathway highlighting the topography in the dorsoventral and mediolateral axis. Each density plot is averaged across the rostrocaudal axis. For each pathway, retrogradely labelled neurons were aligned to the centroid of the CLA_RSP_ pathway (magenta). (**b-d**) Histograms showing the distribution of all retrogradely labelled neurons for each pathway in the dorsoventral (**b**), mediolateral (**c**), and rostrocaudal axis (**d**). (**e**) The neuron counts for each retrogradely labelled pathway across rostrocaudal claustrum locations. (**f)** The proportion of neurons in the dorsal shell, core, and ventral shell, for each pathway. The data for cortical injections into aRSP and pRSP were similar to the RSP labelling (magenta) are shown in **Table 1**, and not plotted here. The number of mice/slices/neurons for each pathway are indicated below. * P < 0.05, ** P < 0.01.

**Figure 6:**
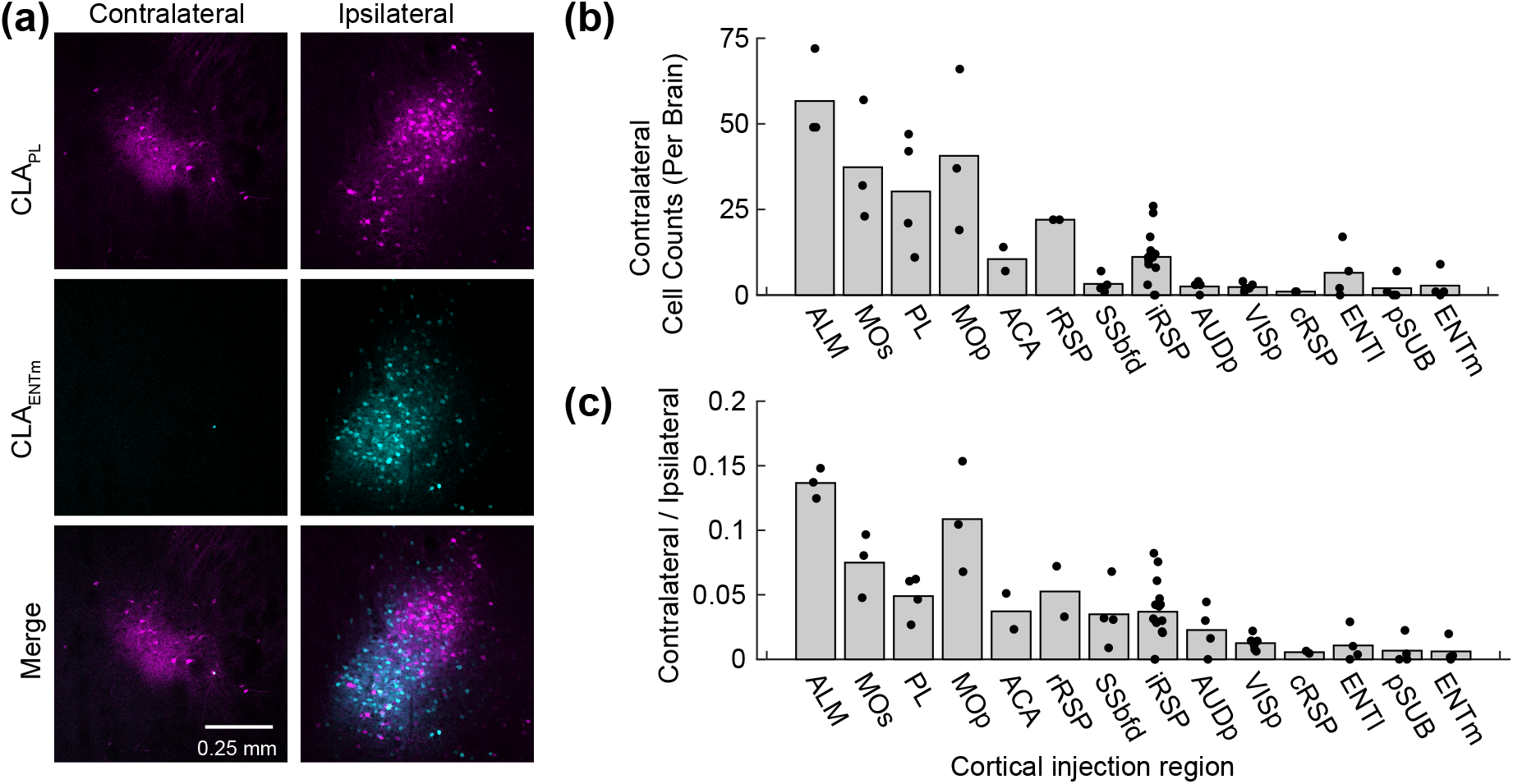
Contralateral projecting claustrum neurons innervate frontal midline cortex but not the temporal lobe. (**a**) A representative experiment showing labelling in the contralateral and ipsilateral claustrum following retrograde tracer injections into PL (magenta, AAV2-retro-GFP) and ENTm (cyan, AAV2-retro-tdtomato). Note the absence of claustrum neurons in the contralateral hemisphere, following retrograde tracer deposited into the ENTm. (**b**) The total number of neurons counted in 5-6 slices in the contralateral hemisphere for each cortical region injection. (**c**) The ratio of contralateral/ipsilateral labelling in the claustrum for each cortical injection region. Contralateral labelled was mainly found in the case of claustrocortical inputs to motor related regions and was less prominent in the case of injections into sensory cortex and temporal lobe. Each point represents one mouse, and the bar plot shows the mean.

Next, we determined the co-projection rate between different claustrocortical pathways (**Figure 7**). With four color tracing, there are theoretically fifteen different labelling patterns that any given neuron can adopt, indicating the projection to one, two, three, or four cortical regions (**Figure 7a-d**). The claustrum co-projection rate was analyzed in twenty-seven pairs of claustrocortical pathways (**Figure 8a-f** and **Table 2**) The vast majority of neurons were only labelled from one pathway (**Figure 8b, d, f, h**). In experiments with four tracers injected along midline spanning ~5mm rostral-caudally (from PL to cRSP), 10% of retrograde labelled neurons were found to project to three or four of the midline regions (**Figure 8f**). With all other injection combinations, the rate of co-projections to all four cortical targets was considerably lower (**Figure 8b, d, h** and **Table 2**). However, co-projecting neurons were common among specific pathways including claustrocortical outputs to ALM/MOp, MOs/RSP, RSP/PL, pSUB/RSP, and pSUB/ENTm, whereas low co-projection rates were found in experiments labelling inputs to sensory cortex (SSbfd, AUDp, and VISp) (**Figure 8a, c, e, g**). The upper limit on the detectability of co-projection between pairs of tracers was found to be ~50-60% (**Figure 9a-f**), suggesting that rates of 10-20% indicate a high rate of co-projection given these methods.

**Figure 7.**
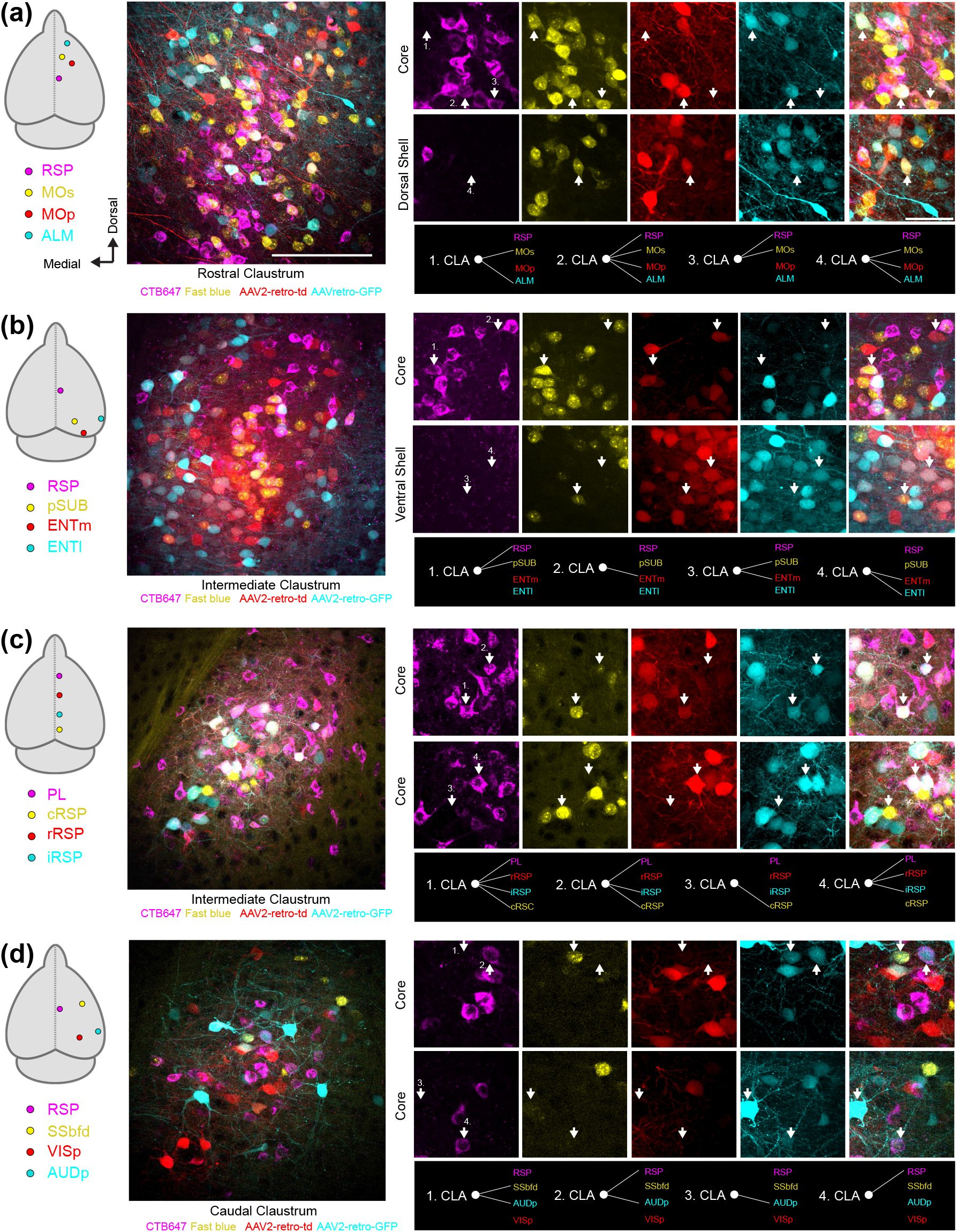
The projection patterns of individual claustrum neurons. (**a-d**) Retrograde tracer injections were made into the regions indicated (far left, as in **Figure 4**) using four different color retrograde tracers. An example field of view from the claustrum is shown with all imaging channels merged. On the right are magnified regions from the larger field of view highlighting examples of the different projection patterns of individual claustrum neurons indicated by white arrows.

**Figure 8.**
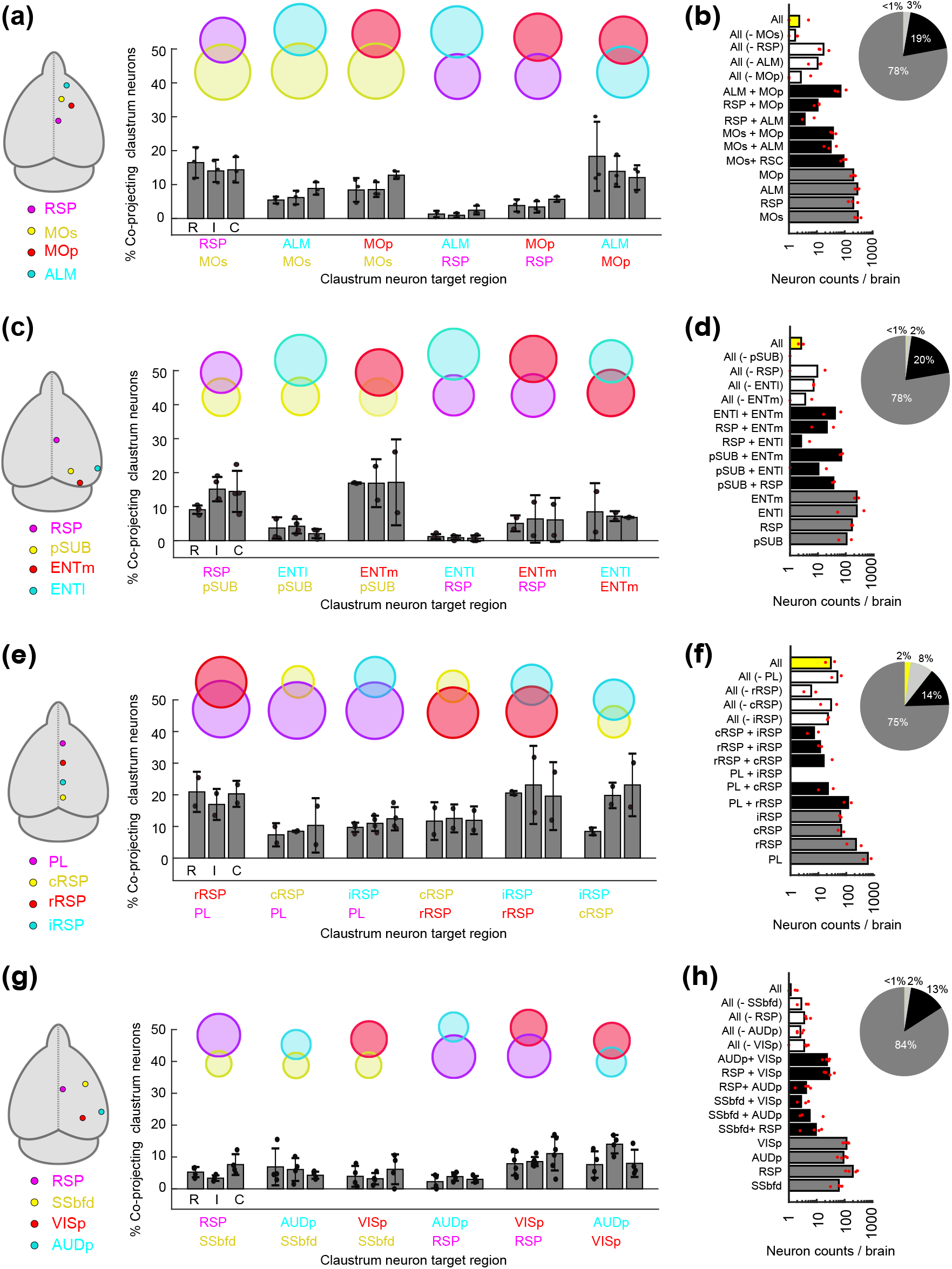
Claustrum neurons project to multiple functionally related brain regions. (**a**) Cortical injection regions, and the percentage of retrogradely labelled neurons projecting to each pair of post-synaptic cortical regions, for experiments with retrograde tracers in RSP, MOs, MOp, and ALM. Each set of bar plots shows the percentage of neurons projecting to both cortical regions as a function of the rostral (R), intermediate (I) and caudal (C) claustrum. The venn diagram is shown above. (**b**) The number of neurons classified into each of fifteen different labelling patterns, ranging from projecting to a single region to projecting to all four regions. The proportion of neurons projecting to one, two, three, or four regions is shown using a pie chart. (**c-d**) The same as A-B but for experiments with retrograde tracers in the RSP, pSUB, ENTm, and ENTl. (**e-f**) The same as (**a**-**b**), for experiments with retrograde tracers in the rostral RSP (rRSP), intermediate RSP (iRSP), caudal RSP (cRSP), and PL. (**g-h**) The same as **a**-**b**, for retrograde tracers in the RSP, SSbfd, AUDp and VISp. Additional experiments with other pathways can be found in **Table 2**.

**Figure 9.**
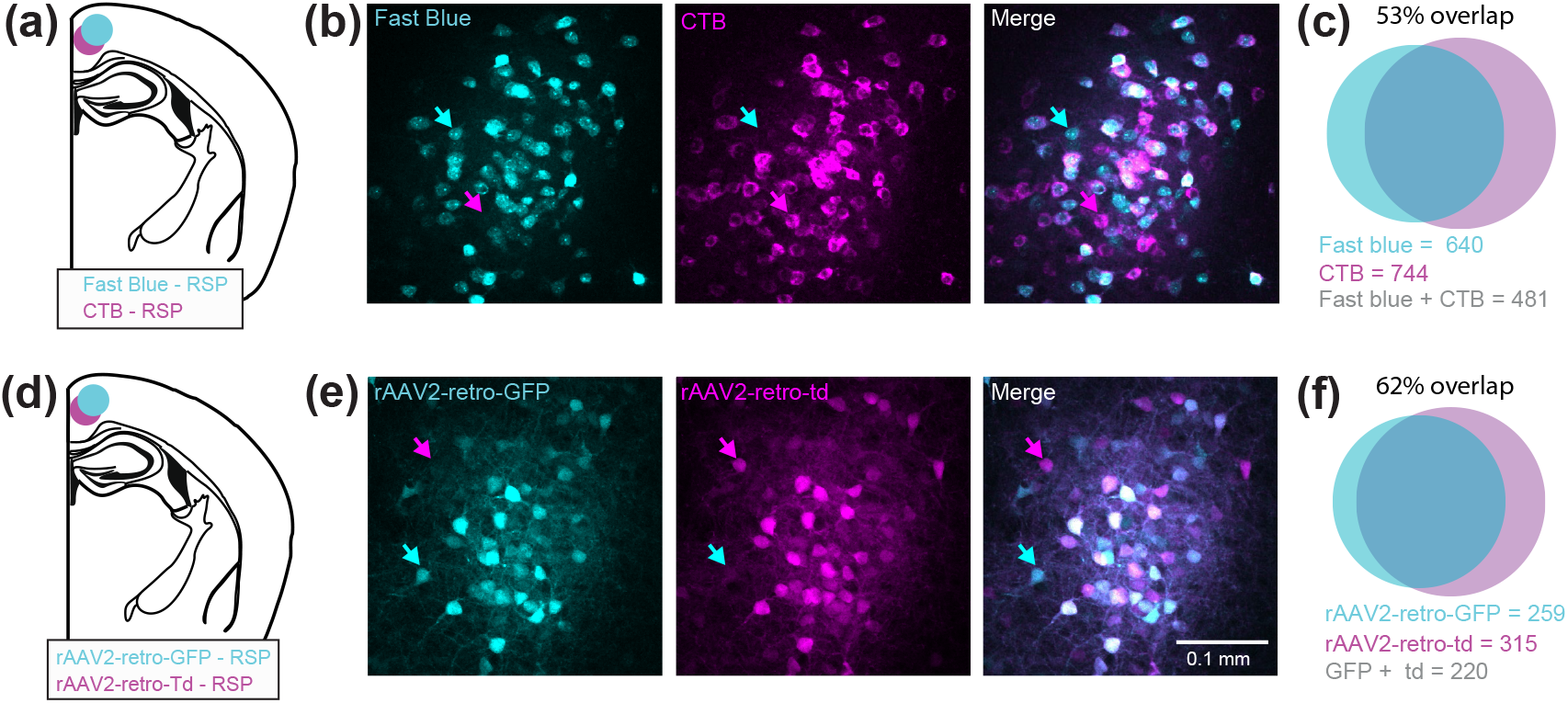
Determining the upper limit of co-projection rates for claustrum projection to the RSP. (**a**) Injections were made to the RSP with pipettes loaded with equal parts Fast blue and CTB647, and a total of 200nl injected. (**b**) Example fluorescence images from the ipsilateral claustrum from one field of view showing a high rate of co-labelling. However, some neurons express Fast blue only (cyan arrow), or CTB only (magenta arrow), indicating that the tracers underestimate the full extent of claustrocortical projections. (**c**) Venn diagram and the number of neurons expressing fast blue, CTB or both, together with the overlap (n = 3 mice). (**d-f)** The same as (**a-c**) but for co-injection of two variants of rAAV2-retro expressing GFP and tdTomato (n = 1 mouse). Note the co-labelling is not complete, showing that each tracer underestimates the extent of CLA_RSP_ projections, and likely other claustrocortical projections as well.

**Table 2:** The number of retrogradely labelled neurons identified in the claustrum for each of the multicolor tracing experiments. The regions where each tracer were injected are indicated in column 3-6, and denoted as A,B,C,D for the remaining columns. The counts shown are uncorrected, meaning that the same neuron can be included in the counts for multiple labelling patterns. For example, a neuron expressing fast blue (A) and CTB (B) would be included in columns counting A, B, and AB. Brains where no data are reported for one channel indicate that the injection site for the particular imaging channel was off target, and the particular channel was not analyzed.

|  |  | Tracer and region |  |  |  | Labelling pattern |  |  |  |  |  |  |  |  |  |  |  |  |  |  |
| --- | --- | --- | --- | --- | --- | --- | --- | --- | --- | --- | --- | --- | --- | --- | --- | --- | --- | --- | --- | --- |
| Mouse ID | Sex | Fast Blue (A) | CTB (B) | rGFP (C) | rTd (D) | A | B | C | D | AB | AC | AD | BC | BD | CD | ABC | ABD | ACD | BCD | AB CD |
| M58 | F | PL | rRSP | cRSP | iRSP | 676 | 667 | 218 | 187 | 255 | 72 | 75 | 120 | 123 | 75 | 59 | 83 | 41 | 68 | 38 |
| M59 | F | cRSP | PL | iRSP | rRSP | 156 | 1012 | 203 | 305 | 87 | 59 | 36 | 99 | 179 | 105 | 41 | 30 | 26 | 84 | 18 |
| M176 | F | MOs | iRSP | ALM | MOp | 482 | 434 | 330 | 280 | 131 | 47 | 62 | 11 | 23 | 60 | 7 | 14 | 14 | 2 | 1 |
| M178 | F | MOs | iRSP | ALM | MOp | 398 | 292 | 393 | 354 | 92 | 36 | 79 | 6 | 33 | 75 | 6 | 19 | 17 | 7 | 5 |
| M231 | M | MOs | iRSP | ALM | MOp | 590 | 255 | 525 | 430 | 98 | 79 | 64 | 12 | 20 | 142 | 2 | 6 | 28 | 3 | 1 |
| M177 | F | SSbfd | iRSP | AUDp | VISp | 119 | 309 | 188 | 176 | 20 | 29 | 9 | 14 | 36 | 37 | 6 | 4 | 8 | 4 | 2 |
| M179 | F | SSbfd | iRSP | AUDp | VISp | 87 | 363 | 109 | 210 | 23 | 6 | 5 | 15 | 51 | 22 | 6 | 3 | 1 | 6 | 1 |
| M180 | F | SSbfd | iRSP | AUDp | VISp | 103 | 146 | 90 | 187 | 12 | 8 | 11 | 6 | 24 | 26 | 1 | 3 | 4 | 0 | 0 |
| M228 | M | SSbfd | iRSP | AUDp | VISp | 65 | 212 | 185 | 182 | 14 | 15 | 17 | 14 | 33 | 46 | 7 | 6 | 7 | 7 | 2 |
| BM12 | M | VISp | iRSP | ACA | - | 141 | 318 | 274 | - | 51 | 44 | - | 74 | - | - | - | - | - | - | - |
| BM13 | M | VISp | iRSP | ACA | - | 263 | 115 | 301 | - | 27 | 81 | - | 29 | - | - | - | - | - | - | - |
| M129 | F | pSUB | iRSP | ENTl | - | 233 | 370 | 689 | - | 79 | 15 | - | 5 | - | - | - | - | - | - | - |
| BM05 | F | pSUB | iRSP | ENTl | - | 330 | 284 | 538 | - | 61 | 31 | - | 5 | - | - | - | - | - | - | - |
| M130 | F | pSUB | iRSP | ENTl | ENTm | 178 | 254 | 73 | 343 | 44 | 6 | 87 | 2 | 46 | 20 | 4 | 10 | 4 | 3 | 3 |
| BM07 | F | pSUB | iRSP | ENTl | ENTm | 312 | 225 | 563 | 450 | 54 | 46 | 89 | 14 | 16 | 87 | 8 | 9 | 20 | 3 | 2 |
| M201 | M | - | iRSP | PL | ENTm | - | 388 | 411 | 325 | - | - | - | 89 | 21 | 27 | - | - | - | 9 | - |
| M205 | M | - | iRSP | PL | ENTm | - | 427 | 347 | 551 | - | - | - | 77 | 28 | 30 | - | - | - | 5 | - |

As suggested by the data in **Figure 8**, the co-projection rate depended on the topography of individual claustrocortical modules (**Figure 10a**). The co-projection rate was positively correlated with the spatial overlap between claustrum modules **(Figure 10b**) and negatively correlated with the distance between downstream cortical targets (**Figure 10c**). Therefore, spatially separated cortical regions, particularly in the rostrocaudal axis, receive input from largely separate sets of claustrum neurons. Finally, we compared the rate of co-projections from the claustrum with the density of corticocortical connectivity. It has been proposed that individual claustrum neurons co-project to anatomically connected regions of the cortex serving to compliment corticocortical connectivity (Jackson et al., 2020; Pearson et al., 1982; Smith et al., 2012; Smith & Alloway, 2014), although this theory has not been rigorously tested. We used previously published corticocortical connectivity estimates (Oh et al., 2014, see methods) to assess the density of anatomical connections between pairs of cortical regions (Methods and Materials). The percentage of co-projecting claustrum neurons shared between claustrocortical pathways was positively correlated with the density of corticocortical connectivity between the two regions (**Figure 10d**). Therefore, subsets of claustrum neurons provide common input to cortical regions with dense interregional connectivity, whereas weakly connected regions receive input from different claustrum outputs.

**Figure 10.**
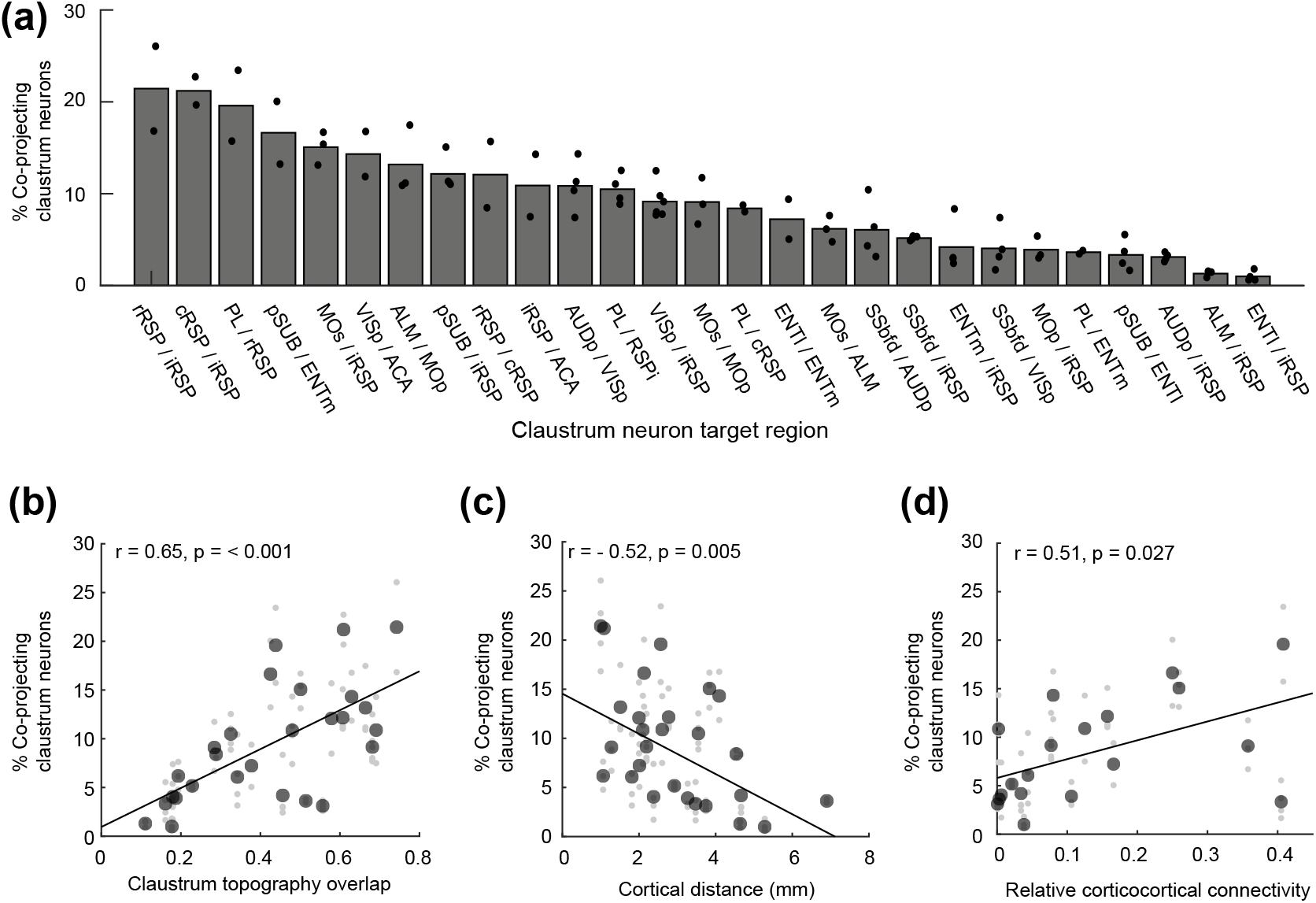
Claustrum projections provide common input to neighboring and connected cortical regions. (**a**) The sorted claustrocortical co-projection rate for all pairs of pathways measured. (**b**) The correlation between the spatial overlap of claustrocortical modules, and the percentage of co-projecting neurons for each pair of claustrocortical pathways. The spatial overlap between claustrum modules was calculated by dividing the area jointly occupied by both pathways by the sum total of both individual pathways. (**c**) The correlation between co-projection rate and the distance between cortical injection sites. (**d**) The correlation between co-projection rate and the average (bidirectional) connectivity between each pair of cortical regions. Cortical connectivity was estimated using the data from the Allen Brain Institute (Oh et al., 2014). The large individual points in (**b-d**) indicate the grand mean across all mice for a given pair of pathways, and small grey points indicate experimental replicates. Correlation coefficients were calculated on the grand mean for each pair of pathways.

Finally, we measured the density of different interneuron subtypes in the claustrum. As the PV neuropil aligns mainly with CLA_RSP_ neurons, many projection neurons in the dorsal and ventral shell reside in low PV neuropil regions. Therefore, other interneuron types may exhibit different topographical arrangements in the claustrum. We determined the density and topography of PV, somatostatin (SST) and neuropeptide Y (NPY) expressing interneurons. The vast majority of SST and NPY cells are inhibitory interneurons (Chittajallu et al., 2013; Xu et al., 2010). SST neurons are well known to provide dendritic inhibition, complementing somatic inhibition provided by PV neurons (Butt et al., 2005; Y. Kawaguchi & Kubota, 1997; Kepecs & Fishell, 2014). SST cell bodies were most dense in the claustrum shell (**Figure 11a-b**), whereas PV cell bodies and neuropil were denser within the core (**Figure 1** and **Figure 11c**). SST neurons were more numerous than PV neurons in the intermediate and caudal claustrum (**Figure 11d-e**), and PV and SST showed inverse patterns of neuropil labelling in the core and shell (**Figure 11f**). Consequently, the spatial profile of SST neuropil labelling was negatively correlated with CLA_RSP_ outputs (**Figure 11g**). NPY cell bodies and neuropil were strongly and uniformly labelled across claustrum core and shell (**Figure 12** and **Figure 13a-d**). In total, the density of NPY (134.4±24.3 cells/ mm^2^) and SST (93.5±18.3 3 cells/ mm^2^) interneurons was greater than PV (43.8±8.23 cells/ mm^2^) (**Figure 13f**). There was a 25% overlap of NPY and SST neurons, particularly in the claustrum shell, but only a 1.6% overlap between PV and NPY (**Figure 13g**). We measured the ratio between different interneuron subtypes in the claustrum and in neighboring brain regions. The SST/PV and NPY/PV ratio was particularly high in the claustrum relative to other brain regions (**Figure 13h**) suggesting an inhibitory neuron signature that aligns more closely to that found in association cortex (Y. Kim et al., 2017). This differential pattern of neuropil and cell body labelling through the claustrum suggests different claustrum output modules are differentially controlled by PV, SST, and NPY mediated inhibition (**Figure 14**).

**Figure 11.**
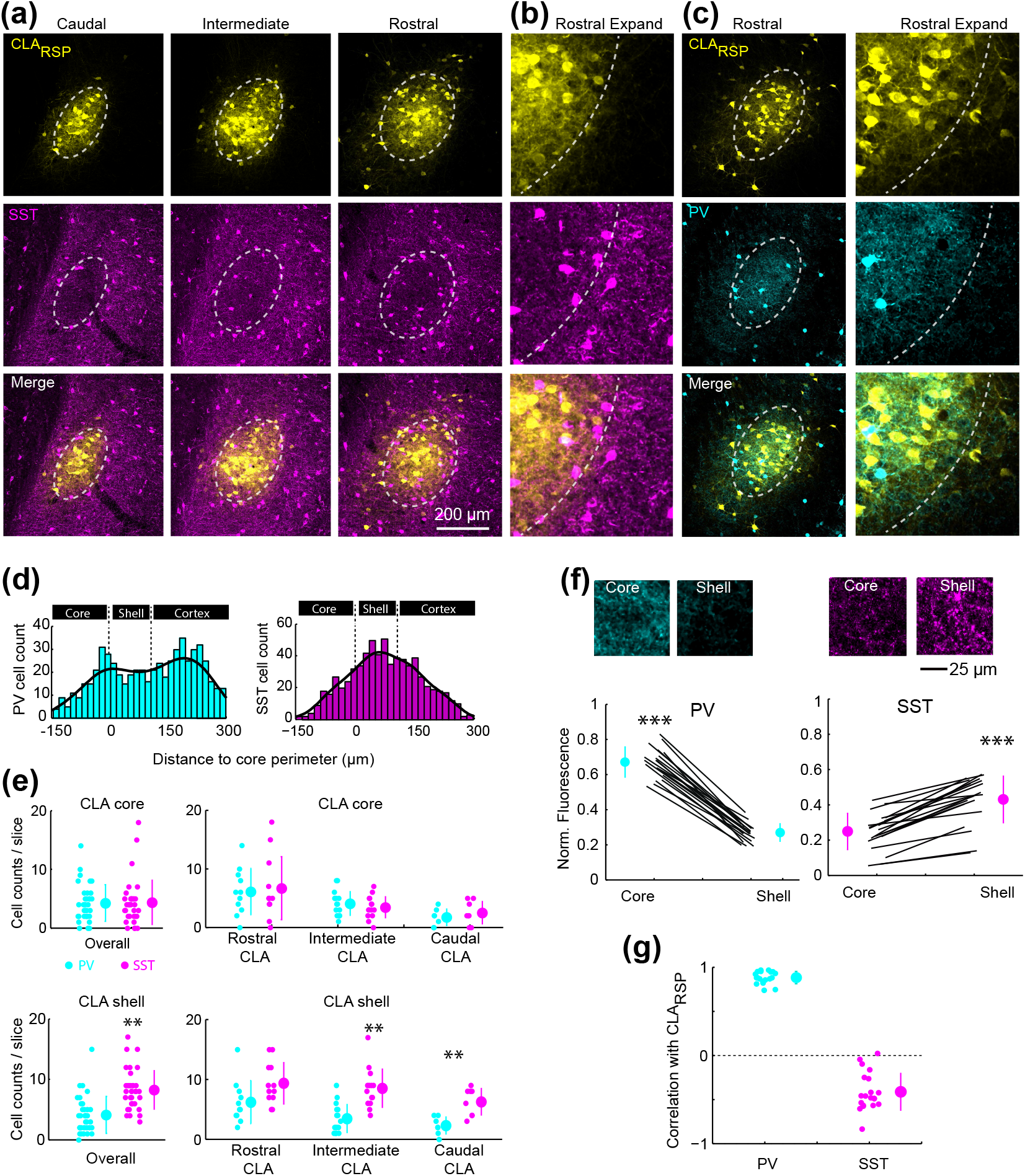
Somatostatin neurons are located in the claustrum shell. (**a**) Example images showing CLA_RSP_ and somatostatin (SST) labelling in the claustrum. (**b**) An expanded view of panel (**a**). (**c**) CLA_RSP_ and PV labelling (as in **Figure 1**) in a brain sections adjacent to panel (**a**). (**d**) The spatial distribution of SST and PV neurons relative to the claustrum core/shell perimeter. (**e**) The number SST and PV neurons in the core and shell of the claustrum across the rostrocaudal axis. The number of PV and SST cells in the core were not different (4.3±3.9 SST cells versus 4.2±3.1 PV cells/slice, t = 0.36, p = 0.72, n= 34 slices from 5 mice). There were more SST cells than PV cells in the shell (8.3±3.3 SST cells versus 4.1±3.1 PV cells/slice, t = 5.52, p = 2.9×10^−6^), and this was mainly due to the PV-SST difference in the intermediate and caudal claustrum. **(f**) The normalized neuropil fluorescence of PV and SST in the core and shell. Example images are shown above. PV neuropil fluorescence was greater in the core (0.67±0.08 versus 0.26±0.05, t =26.9, p = 5.17×10^−16^), whereas SST neuropil was greater in the shell (0.43±0.13 versus 0.25±0.1, t = 10.1, p = 4.3×10^−9^). (**g**) The correlation coefficient between the spatial distribution of CLA_RSP_ and PV (r = 0.88±0.07, n = 19 slices in 3 mice) and SST (r = −0.41±0.21, n = 20 slices in 3 mice). ** P< 0.01, *** P < 0.001.

**Figure 12.**
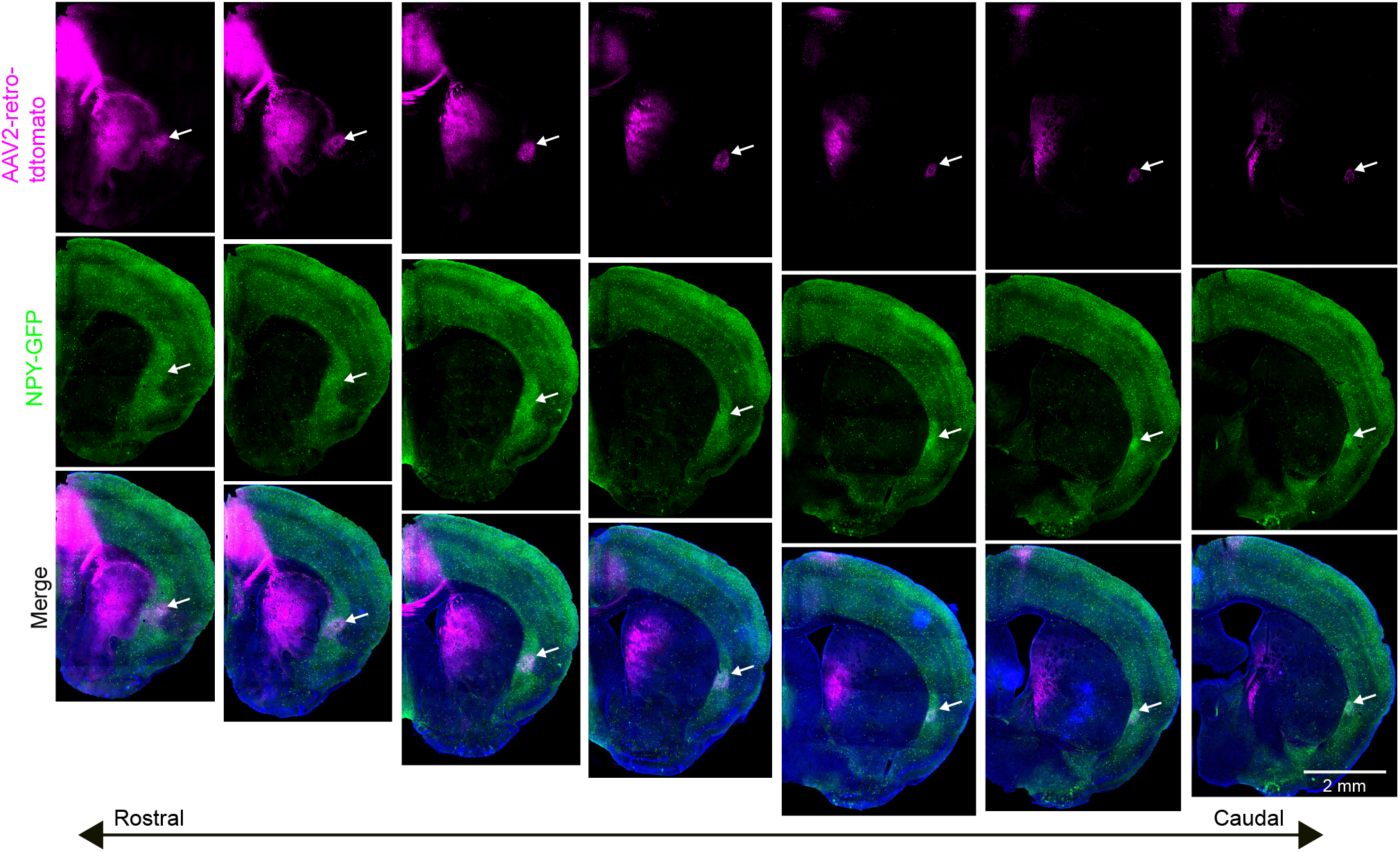
Wide-field imaging of NPY labelling in the claustrum. Wide-field images of coronal sections, sorted according to the rostrocaudal axis (left to right), showing NPY labelling (green) together with retrograde labelling from ACA (magenta). The out of focus excitation with wide-field imaging better highlights the density of NPY neuropil in the claustrum and dorsal endopiriform cortex. The white arrow in each panel indicates the lateral edge of the claustrum. Confocal imaging was performed for all images quantified in the main manuscript.

**Figure 13.**
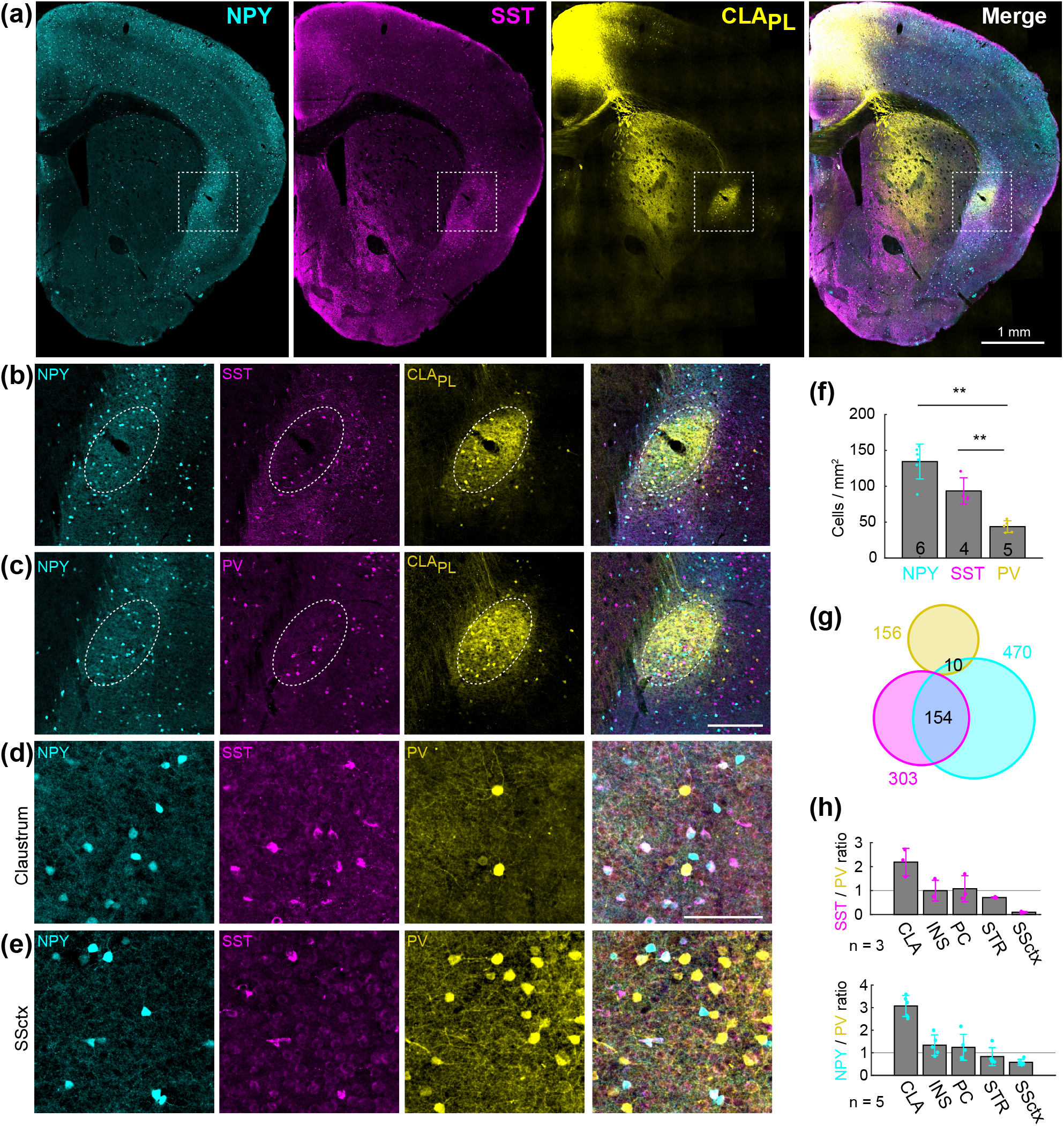
Neuropeptide Y neurons are densely expressed throughout the claustrum. (**a**) Example labelling of neuropeptide Y (NPY), somatostatin (SST), and claustrum - prelimbic cortex (CLA_PL_) neurons. (**b**) Expanded view from panel (**a**), showing the spatial relationship between NPY, SST, and CLA_PL_ neurons. (**c**) As in panel (**b**), but showing NPY, PV, and CLA_PL_ neurons. (**d**) Claustrum immunohistochemical labelling of SST and PV in an NPY-hrGFP mouse. (**e**) As in (**d**), but for the somatosensory cortex of the same slice. (**f**) The density of NPY, SST, and PV neurons in the claustrum. (**g**) Venn diagram showing minimal overlap between interneuron subtypes in the claustrum. (**h**) The SST:PV ratio (top) and NPY:PV (bottom) ratio for the claustrum, insula (INS), piriform cortex (PC), striatum (STR), and somatosensory cortex (SSctx). **P < 0.01.

**Figure 14.**
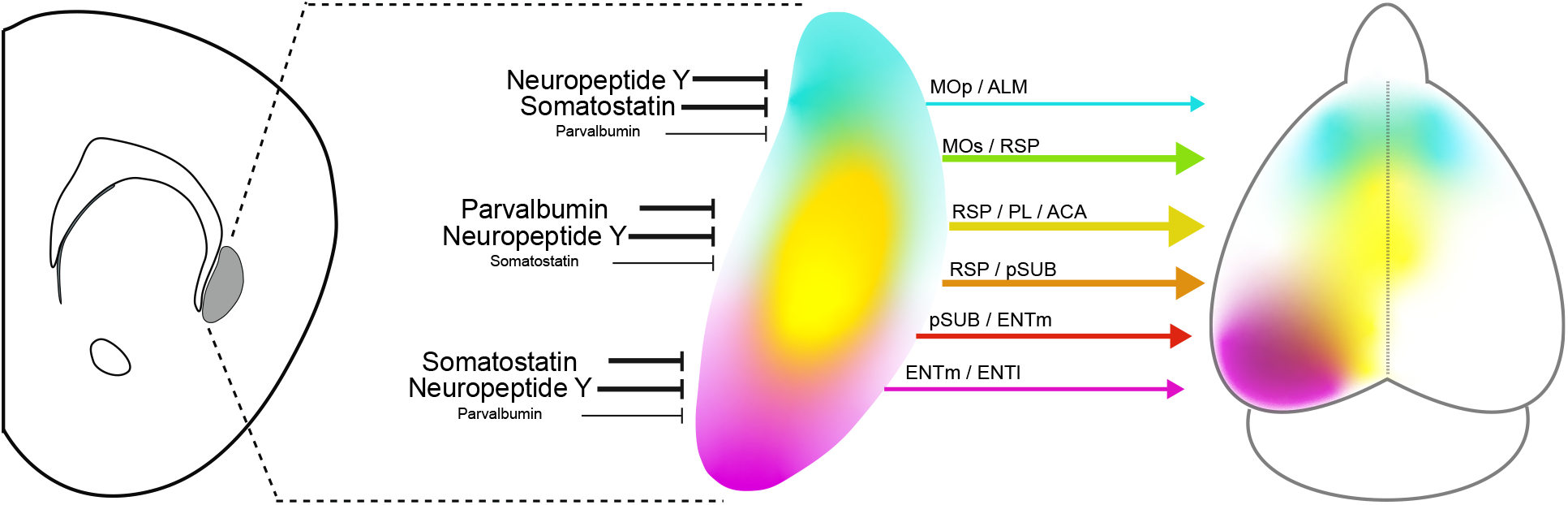
Summary of the topographic mapping between claustrum and cortex. Claustrocortical projections are mainly organized across the dorsoventral axis which maps onto the rostrocaudal axis of the cortex. Interneuron subtypes are differentially localized to the core and shell of the claustrum. The lines linking claustrum and cortex highlight some of the most numerous and divergent claustrocortical pathways identified in this study.

## DISCUSSION

The outputs of the claustrum have been described as both highly divergent or parallel, therefore several details of this system have required resolution. We found that claustrum outputs in the mouse are topographically organized, giving rise to discrete claustrocortical modules which provide common input to neighboring and anatomically connected cortical regions. At the same time, topographically separated modules project to independent cortical regions. Supporting this spatial organization of claustrocortical projections, we found PV, SST, and NPY interneurons each exhibit unique spatial densities and distributions suggesting claustrocortical domains across the dorsoventral axis are controlled by different landscapes of inhibition. These anatomical motifs can support the claustrum coordination of related regions of the cortex while also enabling parallel outputs to participate in different cortical operations.

Previous anatomical investigations into the organization of claustrocortical connections have reported mixed results with respect to topography and axon collateralization. In some cases, different claustrocortical outputs were found to exhibit specific spatial profiles (Kitanishi & Matsuo, 2017; Macchi et al., 1983; Minciacchi et al., 1985; Sadowski et al., 1997; Smith & Alloway, 2014), while in other reports, very little topographical organization was identified (Sloniewski et al., 1986; White et al., 2017). Likewise, claustrum neurons have been reported to co-project to multiple cortical regions (Smith et al., 2012; Y. Wang et al., 2019; Zingg et al., 2018), whereas in other instances little to no co-projections between different claustrocortical pathways were identified (Sloniewski et al., 1986; White et al., 2017).

The discrepancy between studies can be accounted for by several factors. First, claustrocortical mapping has usually focused on a small number of projections in each experiment. Therefore, differences in topography and co-projection rate would depend on the choice of cortical injection target. For example, injections in multiple areas of midline cortex would lead to a high rate of co-projecting neurons and a lack of topographical difference between pathways, whereas injections into temporal lobe and frontal motor areas would lead to low co-projection rate and major topographical differences in claustrum labelling. Our approach involved the study of multiple cortical injection sites and registering the data to a common pathway. This approach has not been used previously, but we find it is essential for accurate registration across experiments where small differences in the location of claustrocortical projection modules arise. Another issue giving rise to discrepancies between studies is the species-specific organization of claustrocortical projections. The original work on claustrocortical connections was performed in cat and primate which show clear topographical zones that project to specific areas of visual, auditory, and somatosensory cortex (LeVay & Sherk, 1981; Olson & Graybiel, 1980; Pearson et al., 1982; Remedios et al., 2010; Witter et al., 1988). However, in rodents, the majority of claustrum neurons project to association cortex, rather than primary sensory cortex (White et al., 2017; White & Mathur, 2018; Zingg et al., 2018). As mice may now provide an essential model system to study the function of the claustrum, the data we present here will enable neural activity of specific claustrocortical pathways to be manipulated or measured while taking into consideration the crosstalk with other projection streams. However, there are species differences in the anatomical organization of the claustrum (Edelstein & Denaro, 2004; Orman et al., 2017; Pham et al., 2019; Smith et al., 2019; Witter et al., 1988) and therefore the results in mice may not generalize to other species.

We found that the claustrocortical system is organized into many independent output pathways, yet it is unclear if each projection stream is comprised of distinct cell types defined by other modalities such as gene expression. Anterograde tracer injections into the claustrum in different transgenic mice show diffuse labelling across anterior, temporal, and midline cortex (Atlan et al., 2018; Narikiyo et al., 2020; Q. Wang et al., 2017; Y. Wang et al., 2019). For example, *Gnb4*-cre mice injected with cre-dependent anterograde AAVs in the claustrum show axons innervating the entire cortical mantle (Narikiyo et al., 2020; Q. Wang et al., 2017). However, upon closer examination using single cell axon reconstruction, it appears that *Gnb4* claustrum neurons can be sub-classified into at least four different clusters (Y. Wang et al., 2019). One cell type sends axons exclusively to the temporal lobe, while another predominantly innervates multiple regions along midline cortex. Therefore, these findings align with our data showing that claustrum neurons typically innervate one set of cortical regions and suggests that *Gnb4* provides genetic access to multiple claustrocortical pathways. A parallel study has shown that claustrum neurons projecting to the ENTl and RSP have different transcriptomic signatures (Erwin et al., 2020, submitted). However, further work will be required to completely dissect the transcriptomic similarity between claustrocortical projections streams.

Similar to claustrum projections, we found that interneurons were also unevenly distributed across the claustrum axes. PV interneurons have been studied anatomically and physiologically in the claustrum (Druga et al., 1993; J. Kim et al., 2016; Mathur et al., 2009; Real et al., 2003; Reynhout & Baizer, 1999) and have been shown to receive cortical input and generate feedforward inhibition onto the claustrocortical neurons projecting to the ACC (J. Kim et al., 2016). However, the density of PV cell bodies and neuropil labelling decreases drastically in both the dorsal and ventral axes implying that claustrocortical neurons projecting to ALM, MOp, ENTm, and ENTl may receive less prominent PV-mediated inhibition. The presence of other interneurons containing calretinin, vasoactive intestinal polypeptide (VIP), NPY, SST, and cholecystokinin have been shown to exist in the claustrum (Graf et al., 2020; Kowiański et al., 2008; Real et al., 2003). Calretinin neurons show a similar spatial distribution to what we describe here for SST (Druga et al., 2015). However, SST and calretinin comprise only a partially overlapping population of ~ 30% (Xu et al., 2006, 2010), and calretinin labelling also includes VIP interneurons which are functionally different than SST cells (Karnani et al., 2016; Y. Kawaguchi & Kubota, 1997; Yasuo Kawaguchi & Kondo, 2002; Pfeffer et al., 2013; Rudy et al., 2011). Therefore, SST labelling reveals a more specific interneuron class, highlighting interneurons which provide dendritic inhibition. NPY interneurons have not been well studied in the claustrum. However, with the NPY-GFP mouse used here, there was dense neuropil and cell body labeling that outlined the claustrum core and shell (**Figures 12 and 13**). A recent study showed that the intrinsic electrical properties of PV, SST, and VIP interneurons in the claustrum were distinct from each other (Graf et al., 2020), similar to cortex. However, to the best of our knowledge, no study has tested or compared the connectivity of SST, NPY, or VIP cells with different claustrocortical connections. Future studies will be critical to test the hypothesis that different output streams are controlled by different inhibitory circuit motifs.

In conclusion, claustrocortical connections are comprised of several overlapping spatial modules arranged in a dorsoventral continuum, topographically aligned with separate cortical networks. Claustrum neurons innervate many functionally related and anatomically connected cortical regions, but claustrum modules projecting to weakly connected and spatially diffuse cortical regions are non-overlapping. This organizational framework may enable distinct behaviors and brain states to be supported by independent claustrum circuits.

## AUTHOR CONTRIBUTIONS

BM designed the project, collected the data, analyzed the data, and wrote the manuscript. AD and RZ collected data, performed immunohistochemistry, and edited the manuscript. JJ designed the project, analyzed the data, wrote the paper, and supervised the project.

## DATA AVAILABILITY

All data are contained within the manuscript, available from the corresponding author upon reasonable request, or accessible at https://osf.io/83ENS/

## ETHICS APPROVAL

All procedures were performed according to the Canadian Council on Animal Care Guidelines and were approved by the University of Alberta Animal Care and Use Committee (AUP2711).

## ACKNOWLEDGEMENTS

We thank Mahesh Karnani, Jeremy Cohen, and members of the Jackson laboratory for comments on the manuscript, the Faculty of Medicine & Dentistry Cell Imaging Center including Xuejun Sun, Steve Ogg, and Greg Plummer for assistance with microscopy, and Albert Lee and Monique Copland for NPY-hrGFP tissue. BM is supported by a studentship from the Neuroscience and Mental Health Institute. JJ is funded by the University of Alberta (Faculty of Medicine & Dentistry, and Department of Physiology), the National Sciences and Engineering Research Council of Canada (RGPIN-2018-05212), the Brain and Behavioral Research Foundation Young Investigator Grant, and Canadian Foundation for Innovation (John R. Evans Leaders Fund), and the Canadian Institutes for Health Research (grant# 426485).

## ABBREVIATIONS

AAV2: Adenoassociated virus 2
AF-647: Alexa Fluor-647
ACA: Anterior cingulate cortex
ALM: Anterior lateral motor cortex
AUDp: Primary auditory cortex
CLA: Claustrum
CTB: Cholera toxin subunit-b
cRSP: Caudal retrosplenial cortex
ENTl: Lateral entorhinal cortex
ENTm: Medial entorhinal cortex
FB: Fast blue
iRSP: Intermediate retrosplenial cortex
MOp: Primary motor cortex
MOs: Secondary motor cortex
NPY: Neuropeptide Y
PBS: Phosphate buffered saline
PFA: Paraformaldehyde
PL: Prelimbic cortex
pSUB: Post subiculum
PV: Parvalbumin
RSP: Retrosplenial cortex
rRSP: Rostral retrosplenial cortex
SSbfd: Somatosensory barrel cortex
SST: Somatostatin
VIP: Vasoactive intestinal polypeptide
VISp: Primary visual cortex

